# Heterotypic intercellular adhesion tunes efficiency of cell-on-cell migration

**DOI:** 10.1101/2025.08.29.673192

**Authors:** Chandrashekar Kuyyamudi, Suhrid Ghosh, Cassandra G. Extavour

## Abstract

Cell migration across epithelial barriers occurs in diverse developmental, immunological, and pathological contexts. Here we investigate the contribution of heterotypic adhesion between migrating cells and epithelial “substrate” cells to transepithelial migration. Using an *in silico* model inspired by the migration of primordial germ cells across the midgut epithelium in the *Drosophila* embryo, we show that heterotypic adhesion modulates migration efficiency in a non-monotonic manner, revealing the existence of an optimal adhesion regime. Consistent with this prediction, *in vivo* overexpression of E-cadherin in germ cells accelerated their exit from the midgut relative to controls. Beyond providing experimentally testable predictions, our model integrates and explains previous observations on the role of heterotypic adhesion in cell-on-cell migration, offering a framework for understanding transepithelial migration across biological contexts.

**Significance Statement:** Cell adhesion is important for cell migration, and when cells migrate on a substrate of other cells (rather than on extracellular matrix), the adhesive properties of both cell types must be considered. However, whether and how dynamic changes in adhesion regulate cell-on-cell migration remains unclear. Here we approach this problem using transepithelial migration of *Drosophila* embryonic germ cells as a case study. We develop an *in silico* model of the migration process that predicts an optimal level of adhesion between migrating cell and substrate cell to achieve efficient migration. *In vivo* live imaging and genetic manipulation experiments uphold the predictions of this model. This suggests that adhesion is not a simple on/off binary parameter regulating migration.

## Introduction

Cells frequently carry out their functions at locations distinct from their site of origin, making the individual cellular motility that is required for cell migration a critical aspect of their behavior (1, 2). To achieve motility, cells require persistent and directed membrane activity, which is frequently guided by external chemical cues (3–6). This directed membrane activity is enabled by the dynamic nature of cell membranes, which exhibit continuous turnover of cytoskeletal components (7–9)

Cell migration, division, differentiation and death enable the development of multicellular organisms from a single cell (10). Cells in various developmental contexts migrate individually as well as collectively (1, 2, 11). Embryonic development unfolds as a tightly regulated cascade of events, where each step must occur within a specific time window to prevent disruption of subsequent stages (12–15). These temporal constraints also apply to cell migration, making not only the path taken but also the timing of arrival critical to proper development (1, 2, 16, 17).

In the developing embryo of the fruit fly *Drosophila melanogaster*, the primordial germ cells (germ cells) are the first cells to form (18). The germ cells form at the posterior tip of the embryo roughly 2.5 hours after egg laying (18, 19). Around 15 minutes after their formation, the embryo undergoes gastrulation, and the cluster of germ cells enters the primordial midgut upon invagination of the surrounding tissue during gastrulation (20). Here, the germ cell cluster eventually disassembles, and the germ cells exit the gut by individually passing through the monolayer of epithelial cells that form the midgut (20, 21). Recent work has implicated juvenile hormones as chemical cues that guide germ cells toward the posteriorly located gonad (22). Complementing this, Wunen/Wunen2, strongly expressed on the ventral side of the midgut, acts as a repellent that drives germ cells to exit the midgut dorsally (23–26). Once outside the midgut, the germ cells continue their migration to join the mesodermal cells that form the somatic gonad primordium (27).

The *Drosophila* adhesion protein E-cadherin is encoded by the *shotgun* (*shg*) gene, and is maternally deposited in the early embryo (28, 29). Germ cells require this maternally deposited E-cadherin both to form a cohesive cluster and to tether to the invaginating midgut primordium during gastrulation (30–32). Regulation of germ cell E-cadherin during internalization into the midgut is orchestrated by the Trapped in endoderm (Tre1) protein, a G protein-coupled receptor (31, 32). Tre1 enables the germ cell cluster to form a radial configuration of polarized cells that detach from the cluster prior to transepithelial migration (31, 32). In embryos laid by females expressing a loss of function *shotgun* allele (*shg*^*A*9–49^), individual germ cells separate from the cluster prematurely (31). Furthermore, in 38% of *shg*^*A*9–49^mutant embryos, germ cells exhibit a delay in crossing the midgut epithelium (31). The posterior midgut, which is the barrier that the germ cells must breach, is made up of epithelial cells (33, 34). Remodeling of the midgut epithelium endows the epithelial barrier with increased permeability, allowing the germ cells to migrate through it (20, 35). High resolution imaging of the germ cells crossing the midgut epithelium has shown that germ cells maintain contacts with the epithelial cells with no evident intercellular space around them (20). In addition, FGF signaling regulates the distribution of zygotic E-cadherin in the posterior midgut to maintain its epithelial integrity (36). FGF mutants display mislocalized E-cadherin and collapse of the midgut lumen, trapping the germ cells within (36).

Germ cells expressing defective E-cadherin (*shg*^*A*9–49^) show delayed transepithelial migration (31), reminiscent of the delay in border cell migration that occurs when nurse cells either lack or overexpress E-cadherin in the *D. melanogaster* egg chamber (37). Together, these observations suggest the existence of a non-monotonic relationship between migration efficiency and adhesion between migrating and substrate cells. We set out to understand the role of heterotypic adhesion between the migrating cells and “substrate” cells in transepithelial migration, taking as a case study the role of intercellular adhesion mediated by E-cadherin in enabling the germ cells to transmigrate. Leveraging genetic tools and two-photon microscopy, we live imaged the process of transepithelial migration *in vivo.* We combined the insights gained from live imaging with previously published observations to formulate an *in silico* model of the process. This model, formulated in the Cellular Potts Model framework (38–40), provides a novel mechanistic understanding of this transepithelial migration process. Our model reveals the existence of optimal heterotypic adhesion between germ cells and the epithelial cells, which maximizes the efficiency of midgut exit. In addition, our *in silico* model yields testable, potentially generalizable predictions about the role of E-cadherin and intercellular adhesion that may be applicable to other systems of heterotypic cell adhesion and migration.

## Results

### I. Germ cells exit the midgut individually and maintain E-cadherin expression throughout exit

To gain insight into the process of germ cell transepithelial migration, we live imaged the process *in vivo* [**Fig. 1b-b”**]. Before migration, the germ cells are contained by the epithelial barrier of the midgut [**Fig. 1b**], and located at a depth of more than 25μ*m* from the cortex within the embryo (41). To visualize the germ cells, we used the previously generated fly line *nos-LifeAct-tdTomato* (42) which specifically labels germ cell membrane and cortex through posteriorly localized maternally deposited protein. The cells in the gut epithelium also show some residual labelling since the gut arises from the invaginating posterior blastoderm. The residual labelling generates contrast and helps us estimate the position of the gut during transepithelial migration.

**Figure 1:**
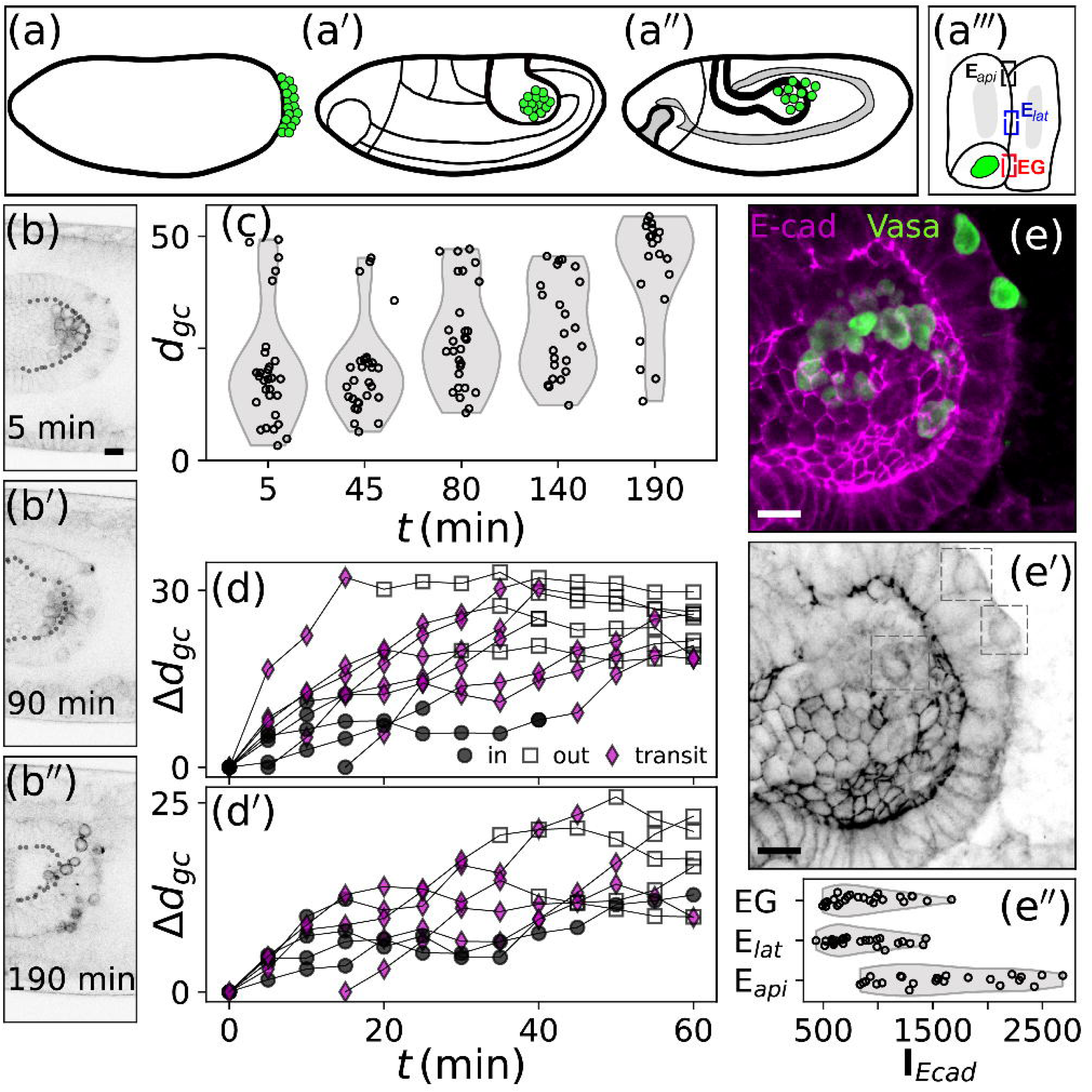
Germ cells exit the midgut individually at comparable speeds. (a-a’’) Schematic of *D. melanogaster* embryonic germ line development. The primordial germ cells are formed at the posterior tip at stage 5. Following gastrulation, they are contained in the midgut at stage 7, and eventually exit the midgut through transepithelial migration at stage 10. **(a’”)** Schematic showing the three different types of intercellular junction where we quantified E-cadherin intensity. **(b-b’’)** Three snapshots from live imaging of the TEM process as the germ cells make their way through the midgut. **(c)** The distribution of the distances of the germ cells from center of the midgut lumen at five different time points are shown. Small numbers under plots indicate sample sizes (number of cells). **(d-d’’)** Movement of germ cells in time measured with respect to their initial (t = 0) position for two different live imaged embryos. Black circles: positions within lumen; magenta rhomboids: positions within the midgut epithelium; white squares: positions outside the midgut. **(e)** Higher magnification of the midgut during germ cell transepithelial migration. Magenta: E-cadherin; green: germ cell-specific protein Vasa. **(e’)** Germ cells express E-cadherin during the transepithelial migration (boxes). **(e’’)** The relative intensity (au: arbitrary units) of the E-cadherin fluorescent signal at the epithelial cell – germ cell junction (*EG*), lateral epithelial – epithelial junction (*E*_𝑙𝑎𝑡_) and apical epithelial – epithelial junctions (*E*_𝑎𝑝𝑖_). n = 24 junctions for each of the three categories. Two-sample Mann-Whitney U tests were used to compute the p values that are shown in panel (e’’) for pairwise comparisons of distributions.

We tracked germ cells throughout transepithelial migration, characterizing their progress at a five minute time resolution [**SI Videos 1 - 3**], and quantified the distances of germ cells from the center to the edge of the midgut lumen as a function of time [**Fig. 1 c**]. **Figure 1d-d’** shows the distance covered relative to their initial position (i.e. t = 0) as a function of time for two different embryos. We found that individual germ cells took 30 minutes on average to go through the midgut epithelial barrier [**Fig. 1d-d’; SI Videos 1 - 3**]. Our live imaging of the transepithelial migration process did not reveal any cell divisions in the midgut epithelial cells that could have generated spaces in the epithelial barrier that would allow for easier exit of germ cells, in contrast to the mechanism that regulates macrophage infiltration in *D. melanogaster* (43).

To examine E-cadherin dynamics during the transepithelial migration, we generated a fly line expressing endogenously GFP tagged Vasa and mScarlet tagged E-cadherin [**SI Methods**]. Using multi-photon confocal microscopy, we observed that E-cadherin in germ cells changed its subcellular localization at the start of the transepithelial migration [**SI Figure 5 a,a’,b, b’; SI Video 6**]. We observed E-cadherin in the epithelia also move laterally upon germ cell entry [**SI Figure 5 c,c’; SI Video 6**]. Imaging individual germ cells at higher time resolution revealed transient E-cadherin foci form at the interface of germ cell and lateral gut epithelia [**SI Figure 5 d,d’; SI Video 7**]. These foci appear to anchor the germ cell to the baso-lateral epithelial membrane as the germ cells moves forward [**SI Video 7**] and exit the gut. Our observations from live imaging were further bolstered by our <u>We also</u> observation of E-cadherin localization on the surface of germ cells during the transepithelial migration, detected by immunostaining for GFP in endogenously GFP-tagged E-cadherin embryos [**Fig. 1e-e’**]. This result is consistent with previously published experiments (31) that detected E-cadherin by immunostaining.

We quantified the intensity of E-cadherin signal at apical and lateral junctions between midgut epithelial cells, as well as at the epithelial-germ cell junctions [**Fig 1e’’**]. Our quantification revealed that the E-cadherin concentration at the germ cell-epithelial cell interface is not significantly different from that observed between epithelial cells on the lateral side (p= 0.31) [**Fig. 1e’’**]. The presence of E-cadherin in germ cells during transepithelial migration, where germ cells physically contact epithelial cells (20), along with the delayed midgut exit previously observed in *shg* loss of function mutants (31), suggests that the expression of E-cadherin in germ cells is functional rather than incidental.

### II. Time to exit the midgut has a non-monotonic dependence on germ cell E-cadherin level

Although inspired by germ cell transepithelial migration, our central question is broader: what role does heterotypic E-cadherin adhesion play in cell migration? To generalize our question beyond germ cell transepithelial migration, we turned to *in silico* modeling, retaining only essential features of the phenomenon. We developed a two-dimensional model of transepithelial migration, where a single motile cell breaches a ring of epithelial cells under the influence of an external, radial chemoattractant gradient [**Fig. 2a**]. Our two-dimensional model, formulated as a Cellular Potts Model (38, 39), enabled us to consider the cells as finite sized objects capable of deformations in shape. We primarily explored the potential role of chemoattractant strength (λ_𝑀_) and the concentration of E-cadherin in the germ cells (𝐶_*G*_) in the time taken for the germ cells to exit the epithelial barrier (τ_𝑒_). We ran our simulations for a maximum of 10,000 Monte Carlo steps, and saved the position of the germ cell as a function of time for each independent iteration for a given choice of system parameters. **Figure 2a** shows a typical snapshot from our CompuCell3D simulation, and **Figure 2b-b’’** shows the modeled germ cell at three different stages of the transepithelial migration. In the CPM framework, the temperature parameter (𝑇) controls boundary pixel activity, where higher temperatures increase the likelihood of energy-raising pixel flips. To investigate how membrane fluctuations at interepithelial junctions affect migration, we modulated the temperature governing pixel acceptance rates at these boundaries. This allowed us to study how epithelial remodeling influences transepithelial migration.

**Figure 2:**
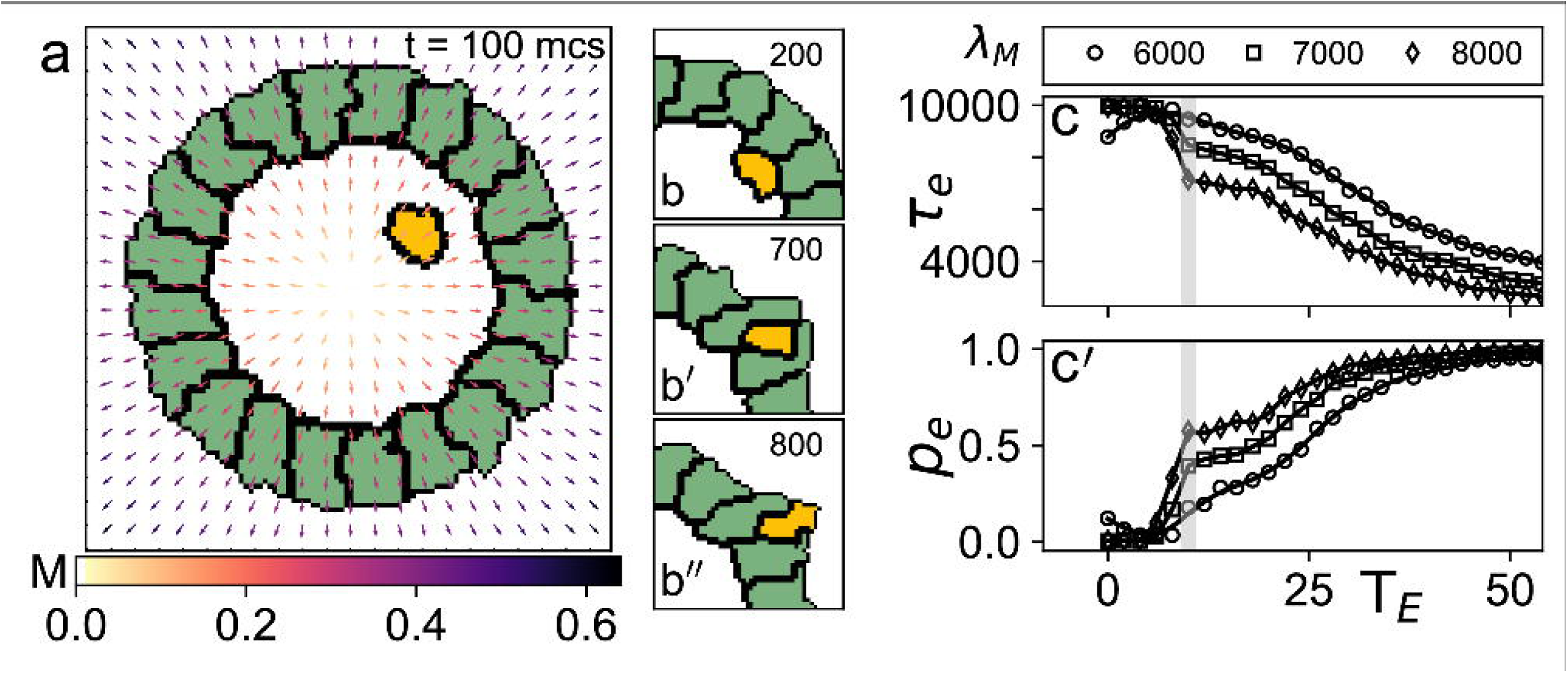
Greater epithelial junctional remodeling aids germ cell transmigration. (**a)** Snapshot of Cellular Potts Model simulation of a germ cell (yellow) exiting the two-dimensional epithelial ring (green). Pixels at the boundary of the cells are shown in black. Colored arrows point in the direction of chemoattractant gradient; colors represent the concentration of the chemoattractant. **(b-b’’)** Snapshots of the germ cell (b) before entering the epithelial barrier, (b’) while inside the barrier and (b’’) just before exiting the barrier. Numbers at top right indicate the time in units of Monte Carlo steps (mcs). **(c)** Average time taken (τ_𝑒_) for the germ cell to exit the midgut decreases with increase in interepithelial dynamics (T_*E*_). **(c’)** Probability of successfully exiting (𝑝_𝑒_) increases with T_*E*_. Trends in c and c’ are independent of the strength of the chemoattractant (λ_𝑀_). The shaded silver line corresponds to 𝑇_*E*_= 10, the value at which al simulations described herein are carried out, unless specified otherwise. Simulation parameters: λ_𝑀_ = 7000, 𝐶_*E*_ = 10, 𝐶_*G*_ = 5, 𝑇 = 10.

Our simulations [**Fig. 2c-c’**] revealed that elevated epithelial temperature increased tissue permeability to germ cells, resulting in faster exit times (𝜏_𝑒_) and higher escape probabilities (𝑝_𝑒_). This is consistent with prior experimental findings where failure in epithelial remodeling of the midgut adversely affected germ cell transepithelial migration (20, 35, 44).

The strength of the modeled chemoattractant 𝜆_𝑀_ beyond which germ cells successfully migrated through the barrier was positively correlated with epithelial cell E-cadherin concentration 𝐶_*E*_ [**Fig. 3a–a’’**]. We interpret this to mean that higher 𝐶_*E*_ values, indicating stronger adhesion between the epithelial cells, offered greater resistance to the trans-migrating cell. Regardless of the value of 𝐶_*E*_, we observed that the time taken to exit the barrier decreased with increased germ cell E-cadherin concentration only up to a point, beyond which there was a sharp increase in the time taken to breach the epithelial barrier [**Fig. 3a–a’’**]. This non-monotonic dependence of τ_𝑒_ on 𝐶_*G*_ suggested that there is an optimal range of heterotypic adhesion between germ cell and the epithelial cell. Within this range, adhesion aids the transepithelial migration, but beyond the range, it hinders germ cell movement. The minima of the curves for different values of λ_𝑀_ in **Figure 3a–a’’** coincide, indicating that the optimal adhesion value for transepithelial migration is independent of the chemoattractant strength. Interestingly, the coefficient of variation in the time to exit τ_𝑒_ as a function of 𝐶_*G*_ has a trend that is inverse of the one exhibited by τ_𝑒_ [**Fig. 3b**]. Specifically, the coefficient of variation is highest when the mean is the lowest, which is not due to standard deviation being independent of 𝐶_*G*_. Instead, the standard deviation increases with 𝐶_*G*_ while the mean decreases with 𝐶_*G*_**[SI Fig. 1]**.

**Figure 3:**
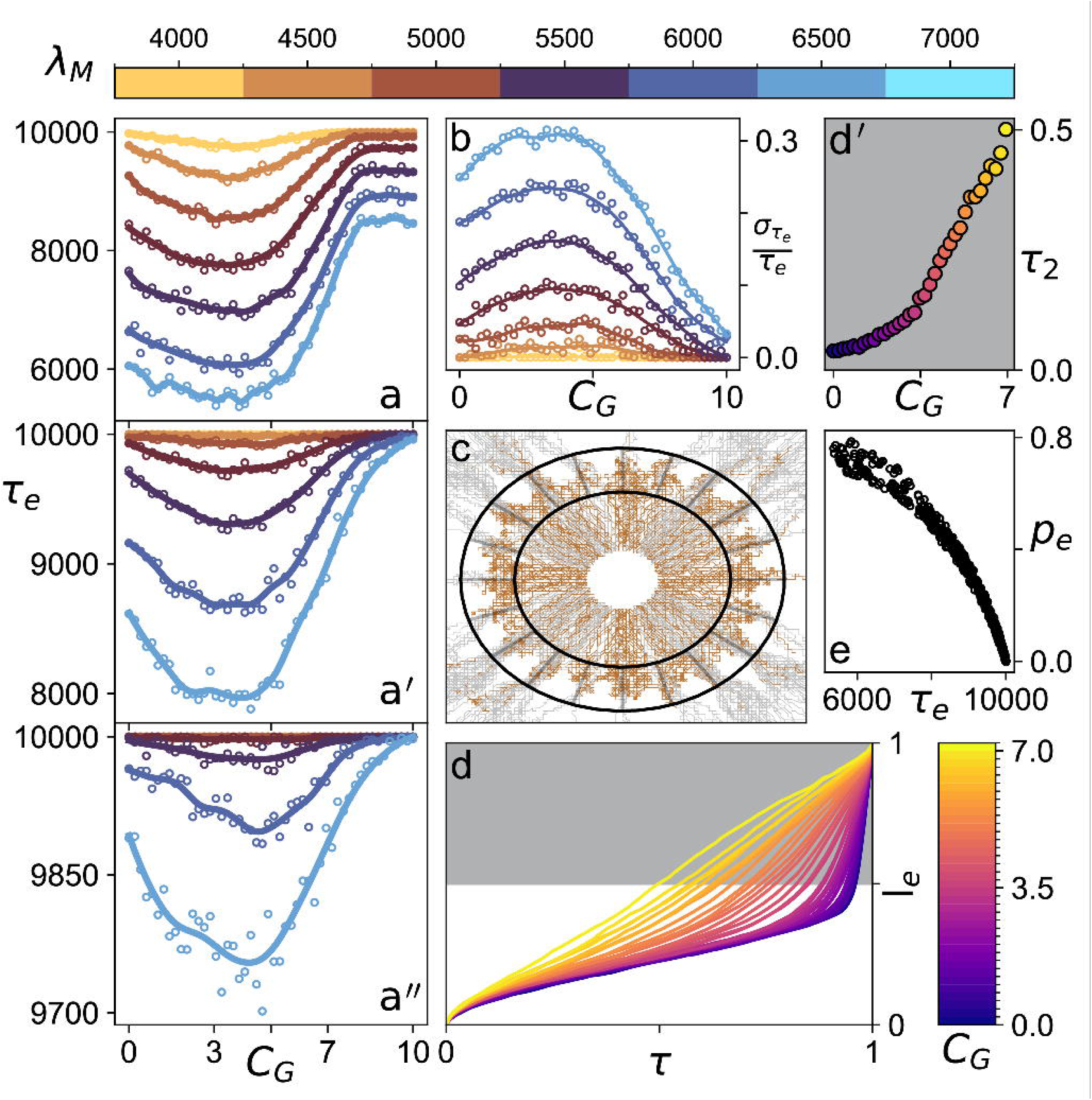
Time to exit the epithelial barrier has a non-monotonic dependence on germ cell E-cadherin concentration. **(a-a’’)** The time a simulated germ cell takes to exit the 2D midgut epithelial barrier as a function of germ cell E-cadherin concentration (𝐶_*G*_). (a), (a’) and (a’’) represent increasing epithelial cell E-cadherin concentrations (𝐶_*E*_ = 8, 𝐶_*E*_ = 10, 𝐶_*E*_ = 12 respectively), offering increasing resistance to the germ cell transepithelial migration. The curves in each of the three panels are colored according to the strength of the chemical field (λ_𝑀_). Regardless of the value of 𝐶_*E*_, for strong enough values of λ_𝑀_ the time to exit shows a minimum that is independent of the λ_𝑀_ value. X-axis label “𝐶_*G*_” underneath **(a”)** applies also to **(a)** and **(a’)**. **(b)** The coefficient of variation in time to exit as a function of 𝐶_*G*_ for the case of 𝐶_*E*_ = 10. **(c)** Trajectories of germ cells for a typical parameter set obtained from 1000 independent stochastic simulations. Silver: successful TEM; brown: germ cell exit failure; black lines: mean position of the epithelial cells forming the 2D barrier. **(d)** Mean distance covered by the germ cell within the barrier (𝑙_𝑒_) as a function of time. The time 𝜏 is normalized by the total time of travel, where 0 represents time of entry into the barrier and 1 represents time of exit. When the germ cell E-cadherin concentration is low, the germ cell covers much of the distance in a very short interval of time at the end of the journey. Higher germ cell E-cadherin levels lead to the germ cell migrating at roughly uniform speed throughout the journey. **(d’)** The time taken to cover the second half of the journey 𝜏_2_ as a function of germ cell E-cadherin concentration. **(e)** The relationship between 𝜏_𝑒𝑥𝑖𝑡_ and 𝑝_𝑒𝑥𝑖𝑡_ is shown using data aggregated by simulating the model over various values of λ_𝑀_, 𝐶_*G*_ and 𝐶_*E*_. 𝜏_𝑒𝑥𝑖𝑡_ decreases monotonically with 𝑝_𝑒𝑥𝑖𝑡_, implying that the shorter the time to exit, the higher the chances of exiting.

To better understand the potential role of heterotypic adhesion in determining the speed of exit, we focused on the trajectories of the migrating cells at different values of 𝐶_*G*_. We ran our simulations for a maximum duration of 10,000 Monte Carlo steps (45), and the cells which had not exited by then were considered to have failed to exit. In **Figure 3c**, the trajectories from ∼1000 independent iterations are shown for 𝐶_*G*_ = 4.89, 𝐶_*E*_ = 10.0 and λ_𝑀_ = 7000. While there are numerous instances of the migrating cell failing at early stages of the transepithelial migration, we see many instances of germ cells stuck to the outer edge of the epithelial barrier and failing to complete midgut exit. At low values of CG, germ cells are stuck closer to the apical side of the midgut, whereas at high values germ cells get attached to the basal side, with much of their cell body outside the barrier [**SI Figure 2**]. We considered all the successful trajectories for 𝐶_*E*_ = 10 and λ_𝑀_ = 7000 and normalized the time for each trajectory, such that τ = 0 and τ = 1 mark the start and end of the transepithelial migration respectively. The time-normalized mean trajectories are shown in **Figure 3d**, colored according to their values of 𝐶_*G*_. We observed that germ cells with lower E-cadherin concentrations spent much of their time getting to the middle of the epithelial barrier, while the second half of the journey, culminating in complete exit from the midgut, was quite rapid. The fraction of total time spent in getting to the middle dropped with increasing germ cell–epithelial cell adhesion, with speed through the barrier becoming constant for high values of 𝐶_*G*_ (∼7.0). To highlight the difference in the fraction of total travel time spent at different halves of the epithelial barrier, we show the fraction of time spent to cover the second half of the epithelial barrier (referred to as τ_2_) in **Figure 3d’**. τ_2_ showed a nonlinear increase with 𝐶_*G*_ approaching 0.5 at lager values of 𝐶_*G*_. The probability of success in exiting the epithelial barrier had an inverse non-linear relation to time to exit τ_𝑒_ regardless of the 𝐶_*E*_, 𝐶_*G*_ and λ_𝑀_, as evident from the scatter plot in **Figure 3e**. The model also predicts that there is an optimal concentration of E-cadherin in the substrate [**SI Figure 4**], consistent with the hypothesis that the efficiency of transepithelial migration is a function of the heterotypic adhesive interface, rather than of the migrating germ cells alone.

In accordance with previous genetic studies (25, 46, 47), we hypothesized that Wunen/Wunen2, which is strongly expressed ventrally, repels germ cells and biases their exit toward the dorsal side. When we incorporated a dorsoventral chemical gradient alongside the radial gradient, germ cells exited the epithelial barrier more dorsally, supporting this hypothesis [**SI Figure 3**].

### III. Overexpression of E-cadherin in germ cells leads to faster exit through the midgut

The results of our model suggested that increasing E-cadherin levels in germ cells would enable them to complete the most time-consuming part of their journey, namely traversing from the middle of the midgut lumen through the luminal epithelial surface, more rapidly, thus resulting in a faster net transepithelial migration time. To test this prediction *in vivo*, we employed the UAS-GAL4 system (48). Maternally deposited E-cadherin serves as the primary source of E-cadherin for all early embryonic cells, including both germ cells and somatic cells (29, 31, 49). Notably, even during transepithelial migration (∼3 hours after egg laying), germ cell E-cadherin remains predominantly of maternal origin, making RNAi-mediated knockdown in germ cells impractical without destroying embryonic integrity (29, 31, 49).

Given these constraints, to assess the effects of E-cadherin overexpression on transepithelial migration, we analyzed germ cell progression in embryos of four different maternal genotypes, collected within 30 minutes from synchronously ovipositing mothers and aged for five hours, as follows: (I) **WT**: Outcrossed wild-type control, expressing endogenous levels of E-cadherin in all cells [*nos-Gal4/+*]; (II) **Pan-Ecad-OE-A**: with ubiquitous E-cadherin overexpression in all cells [*nos>Ecad*]; (III) **Pan-Ecad-OE-B**: with ubiquitous GFP-tagged E-cadherin overexpression in all cells [*nos>Ecad-GFP*]; and (IV) **GC-Ecad-OE**: with germ cell-targeted overexpression of mClover2-tagged E-cadherin [*nos> Ecad-mClover2-nosTCE-pgc 3′UTR*]. To assess the extent of overexpression achieved, we quantified germ cell E-cadherin relative to epithelial apical and lateral expression using anti-DE-cadherin immunostaining [**SI Figure 7**]. The results of this quantification confirmed that use of the ubiquitous driver increased E-cadherin levels in both germ cells and the somatic epithelium, and that the germ cell driver increased E-cadherin levels in germ cells but not somatic cells [**SI Figure 7**]. The position of germ cells relative to the midgut lumen boundary in representative embryos from each group is shown in **Figure 4a–a’’’**. Quantitative analysis revealed that germ cells in all overexpression lines (II–IV) were located farther from the lumen center compared to controls (I) [**Fig. 4b**]. Furthermore, both the mean germ cell distance from the lumen center [**Fig. 4c**] and the fraction of germ cells that had exited the midgut [𝑓_𝑒𝑥𝑖𝑡_ in **Fig. 4d**] were higher in overexpression groups, with the most pronounced effect in the germ cell-targeted overexpression line (IV).

**Figure 4:**
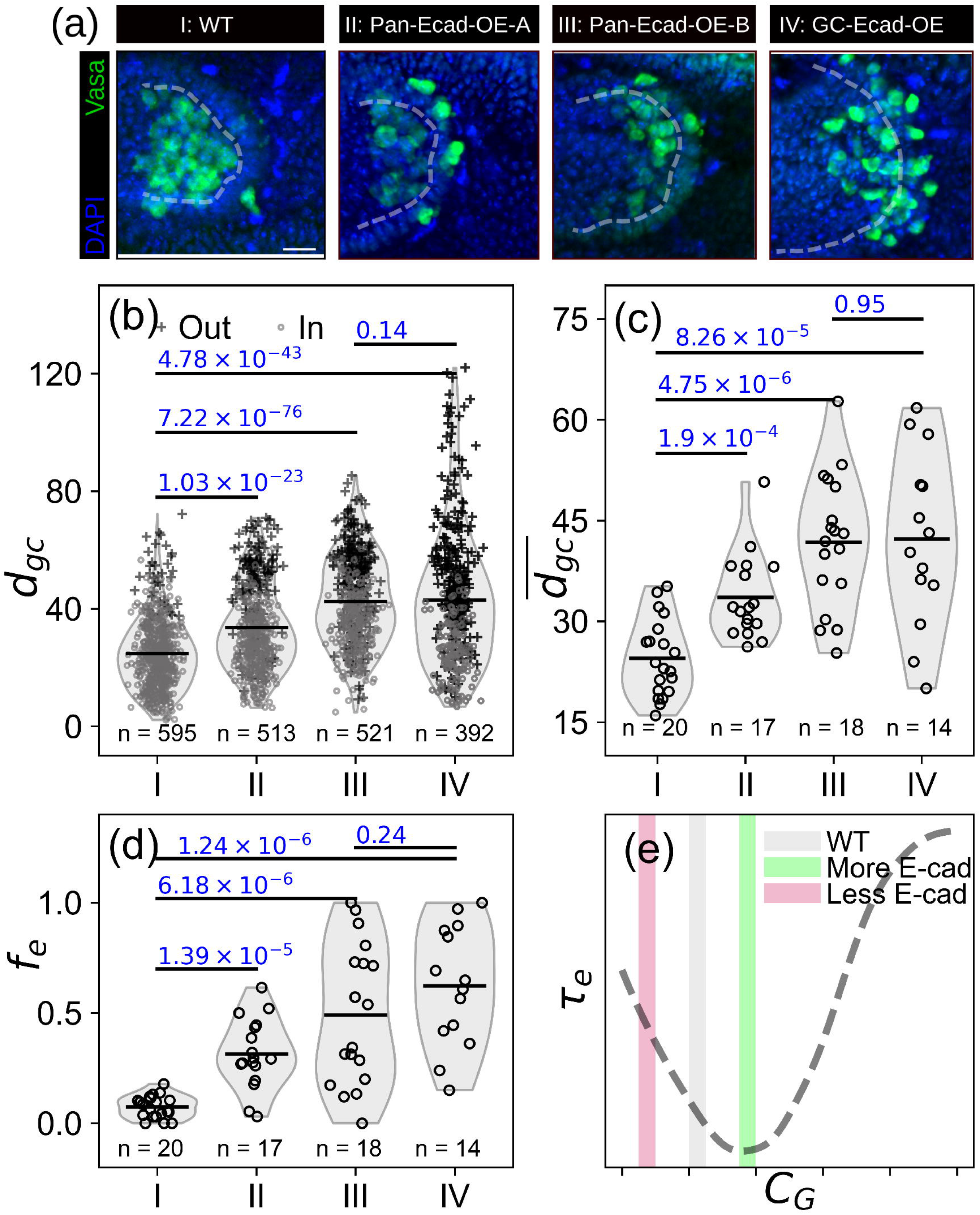
Higher germ cell E-cadherin promotes quicker midgut exit *in vivo*. **(a)** (L-R) Representative micrographs showing the extent of transepithelial migration in the four experimental groups. I corresponds to the control, II and III represent lines where E-cadherin is overexpressed in all cells and IV corresponds to embryos where E-cadherin overexpression is targeted to the germ cells. **(b)** The distribution of distances of germ cells from the center of the lumen for the four experimental groups. Gray and black symbols represent cells inside and outside the gut respectively. The distances of the germ cells from the lumen are largest in the condition with increased germ cell-targeted E-cadherin overexpression (IV). Small numbers under plots indicate sample sizes (number of germ cells). **(c)** Mean distance of the germ cells from the center of the lumen in each embryo for the four experimental groups. Small numbers under plots indicate sample sizes (number of embryos). **(d)** Fractions of germ cells per embryo that have exited the midgut by 5.25 hours after egg laying in the four experimental groups. Small numbers under plots indicate sample sizes (number of embryos). Two sample Mann-Whitney U test was used to compute the p values that are shown in panels **(b)-(d)** for pairwise distribution comparisons. **(e)** We hypothesize that the WT embryos (group I) have 𝐶_*G*_ in the pink shaded region, and that germ cell-targeted E-cadherin expression pushes the system towards the green shaded region, leading to shortening of 𝜏_𝑒_.

These results suggest that higher levels of E-cadherin in germ cells accelerates transepithelial migration, consistent with our theoretical model. We favor the interpretation that wild-type germ cell E-cadherin expression is associated with an intermediate time to exit [gray bar, **Fig. 4e**]. Under this interpretation, loss of function *shg* mutants like *shg*^*A*9–49^(31) would drive the embryo toward the red region shown in **Figure 4e**, associated with longer τ_𝑒_, while *shg* overexpression would drive it towards the green region in **Figure 4e** associated with faster germ cell exit.

Our model predicted that elevated epithelial E-cadherin would hinder migration by increasing resistance to germ cell exit. Germ cells in our experiments overexpressing E-cadherin in both somatic and germ cells showed reduced germ cell exit times. This apparent discrepancy can be explained by the hypothesis that wild-type E-cadherin levels in somatic epithelial cells are already near saturation. Under this interpretation, finite adhesion junction availability and membrane transport limitations would prevent additional E-cadherin from forming new junctions at high concentrations (50–53). Thus, in embryos with E-cadherin overexpression in both germ cells and somatic cells, like the ones we examined herein (**Fig. 4**, groups II and III), we hypothesize that E-cadherin levels higher than wild type levels would have a stronger effect on germ cells than on epithelial cells, effectively mimicking germ cell-specific overexpression.

To further test this, we drove zygotic expression of GFP-tagged E-cadherin specifically in the gut using the 48Y-GAL4 endodermal driver (54) [**SI Figure 6a, b**]. We observed only a small, statistically insignificant decrease in the fraction of germ cells that had exited the gut relative to controls [**SI Figure 6f**]. Because the 48Y-GAL4 driver drives detectable expression 30 minutes after transepithelial migration has initiated [**SI Figure 6b**], it could be the case that our experiment changed E-cadherin levels too late to severely perturb the process. However, this result suggests that overexpression of E-cadherin in the already E-cadherin-rich epithelium has little effect on transepithelial migration, in contrast to germ cell-specific overexpression.

We next used the 48Y-GAL4 driver to induce gut-specific knockdown of E-cadherin (shg-RNAi) and observed a statistically significant reduction in the fraction of germ cells exiting the midgut compared to wild type controls [**SI Figure 6f**]. This phenotype mirrors what Parés and Ricardo (36) reported for FGF pathway mutants, where mislocalization of epithelial E-cadherin caused a substantial proportion of germ cells to fail to exit the midgut. These observations are consistent with our hypothesis that epithelial E-cadherin, and not just germ cell E-cadherin, is required for successful germ cell transmigration. We note that we cannot rule out the alternative hypothesis that germ cell transepithelial migration is impeded in this condition because of broad defects in gut architecture that might be present since epithelial E-cadherin is integral to the structural integrity of the gut and its lumen (36). However, if the gut epithelium were indeed weaker because of E-cadherin depletion, we would expect more efficient germ cell exit, which is the opposite of what we observed. We therefore favor the hypothesis that E-cadherin expression in the gut epithelium is an important contributor to the heterotypic interactions that are required for germ cell exit from the gut.

## Discussion

Our work sheds light on the potential role of heterotypic E-cadherin adhesion during the transepithelial migration of germ cells through the midgut. Further, the predictions of our *in silico* model could in principle apply to any heterotypic adhesive migration process, as we incorporated only a few features specific to germ cell transepithelial migration. Our model logic consists of three key elements, i) a migrating cell that forms transient heterotypic adhesive contacts with a barrier epithelium/endothelium, ii) a chemoattractant gradient providing directed motility, and iii) an optimal adhesion range, where too little adhesion fails to provide the traction needed to breach the barrier, while too much adhesion impedes forward movement by making junction disassembly rate-limiting. We believe this logic is general, while the migratory and substrate cell type, the molecular identity of the adhesion molecules and the source of the driving force are specific to the system.

Our results suggest a non-monotonic dependence of migration efficiency on the heterotypic adhesion between the migrating cell and the substrate cells [**Fig. 4d**]. We posit that traction is essential for cell migration, and that transmembrane proteins like integrins facilitate traction by anchoring cells to the extracellular matrix, thereby enabling forward movement (55). Additionally, cadherin-like molecules mediate homotypic adhesion, allowing cells of the same type to form cohesive tissues such as epithelia (49, 56). These molecules also participate in heterotypic adhesion when migrating cells interact with others expressing cadherins (57–59). A notable example is the transendothelial migration of melanoma cells, where heterotypic adhesion is clearly observed (57). Melanoma cells form heterotypic N-cadherin mediated adhesion junctions with the endothelial cells during their migration (57, 60). Based on our model, we predict a non-monotonic relationship between transmigration efficiency and the heterotypic adhesion between melanoma cells and endothelial cells. More broadly, diapedesis - the process by which cells breach endothelial barriers - relies on heterotypic interactions between transmembrane adhesion proteins (58, 59). The prevalence of heterotypic adhesion during transepithelial migration suggests that our work may have broader implications beyond the specific context of *Drosophila* primordial germ cell migration.

Many conserved features of cell migration are applicable well beyond a specific biological context. In a migrating cell, actin filaments polymerize at the leading edge and undergo retrograde flow, functioning much like the rotation of wheels in a car (61–63). However, the forward motion of a cell depends on its physical coupling to the substrate, just as the tires of a car grip the road to provide traction (61, 62, 64). This mechanism is described by the *molecular clutch model* of cell motility (65). Adhesion molecules such as integrins and cadherins allow cells to bind to surfaces or to other cells, engaging the molecular clutch (65, 66). Integrin-mediated traction is typically used when cells migrate over an extracellular matrix, whereas cadherins facilitate cell-on-cell migration, as seen in the case of border cell migration (37, 66), and in the transepithelial migration case that we consider here.

Our *in silico* model predicts that an intermediate adhesion strength between migrating cells and substrate cells optimizes transmigration. Our finding aligns with previous observations on *D. melanogaster* border cell migration in the egg chamber (37, 67). In the case of border cell migration, a cluster of six to eight border cells, enclosing a pair of non-motile polar cells, migrates through a “substrate” formed by nurse cells (67). Of these three cell types, polar cells express the highest levels of E-cadherin, which they use to anchor to motile border cells, which have the second highest E-cadherin levels (37). Meanwhile, border cells form transient junctions with nurse cell E-cadherin (which is at the lowest levels of the three cell types in the system), pulling on nurse cells as they migrate (37). Notably, E-cadherin knockdown in nurse cells impedes migration more severely than knockdown in border cells, whereas E-cadherin overexpression in nurse cells slows migration (37). These observations further support the concept that intermediate heterotypic adhesion is optimal for transmigration, specifically E-cadherin enabling the engagement of a molecular clutch (66).

Since the Cellular Potts model framework is rooted in Steinberg’s differential adhesion hypothesis (38, 39, 68), transepithelial migration can be interpreted through this lens. In this framework, epithelial cells, by virtue of their high E-cadherin expression, strongly prefer to bind one another, creating a thermodynamic barrier that the germ cells need to breach to intercalate into the epithelium. Germ cell expression of E-cadherin allows them to compete directly with these homotypic epithelial junctions. While germ cells form fewer adhesive contacts than two epithelial cells can form with each other, their E-cadherin expression is sufficient to make heterotypic binding energetically accessible. The chemotactic gradient then provides the directional bias that tips the overall energy balance in favor of heterotypic adhesion. Migration in the absence of sufficient intercellular adhesion resembles climbing a slippery ladder— lacking the traction needed for upward movement. Conversely, excessive adhesion is akin to ascending a ladder coated in glue, where detachment becomes the limiting factor. Thus, an optimal level of “stickiness” is crucial for efficient migration. Inherited maternal content is known to influence germ cells’ ability to migrate successfully to the gonad (26). A component of this maternal content is the E-cadherin used during transepithelial migration is one component of this maternally supplied content. We speculate that the level of E-cadherin in each germ cell may contribute to germ cell fitness during their journey to the gonad.

## Methods

### Fly stocks and maintenance

We generated an endogenous C-terminal *shotgun*–mScarlet (referred to as E-cadherin-mScarlet) fusion using scarless CRISPR/Cas9-mediated homology-directed repair in *Drosophila melanogaster* (69), using homology arms flanking the *shotgun* stop codon. We identified HDR events by 3xP3-DsRed fluorescence and validated correct targeting by PCR [**Supporting Methods**]. Fly lines *w[1118]; P{w[+mC]=GAL4::VP16-nanos.UTR}CG6325[MVD1]* (abbreviated herein as *nanos-gal4-VP16*; stock #4937), *w[1118]; P{w[+mC]=UASp-shg.GFP}5B* (#58445), *y[1] w[*]; TI{TI}shg[GFP]* (#60584), and *w[*]; P{w[+mC]=UASp-shg.R}5* (#58494) were obtained from the Bloomington Drosophila Stock Center (Indiana, U.S.A.). Fly line *w[*]; TI{TI}vas^EGFP.KI^*(Vasa-GFP; stock #118616;) was obtained from the Kyoto Stock Center (Kyoto, Japan). Fly lines *w; nos-Lifeact-tdTomato-P2A-tdKatushka2-CAAX (Lifeact-tdTomato* landing site VK00027; abbreviated herein as *nos-LifeAct-tdTomato*) and *w; UASp-DE-cadherin-mClover2-nosTCE-pgc 3′UTR* (landing site attP2) *(Lin et al., 2022)* were a gift from Ruth Lehmann (Whitehead Institute, MIT, USA). All flies and crosses were maintained on standard fly medium (0.8% agar, 2.75% yeast, 5.2% cornmeal, 11% dextrose) in an incubator at 25°C, 65% RH and 12H:12H light-dark cycle.

### Fly crosses and embryo staging

To generate over-expression of E-cadherin during transepithelial migration, UASp (over-expression) or Oregon R (wild type control) lines were crossed to *nanos-gal4-VP16* and heterozygous F1 virgin females were collected. 40-50 F1 virgin females were then crossed with 8-10 Oregon R males and transferred into an egg collection cage with a 5mm apple juice agar plate as base (70). The *nanos-gal4-VP16* line is highly expressed in the maternal germ line and deposited in the embryo but is not zygotically expressed in the germ cells or somatic cells during TEM. In the case of *nos-LifeAct-tdTomato* and *Vasa-GFP, E-cadherin-mScarlet* flies, an approximate 1:1 ratio of homozygous males and females were transferred into an identical cage setup. Egg laying was allowed to proceed in 30-minute windows and embryos were aged for five hours after the midpoint of the egg laying window. After collection, embryos were either fixed (embryonic over-expression) or used for live imaging (*nos-LifeAct-tdTomato* and *Vasa-GFP, E-cadherin-mScarlet)*.

### Live imaging

Appropriately staged embryos were manually dechorionated with forceps on double-sided scotch tape and attached to a standard 90 mm polystyrene Petri dish using heptane glue. The dish was then filled with sterile distilled water and mounted on a Zeiss LSM 980 multi-photon microscope. Using a 20x water immersion objective, the total volume of the embryonic gut was imaged at a two-photon excitation wavelength of 1000nm (*nos-LifeAct-tdTomato)*, and 900nm/1050nm (*Vasa-GFP, E-cadherin-mScarlet*), every 1, 5 or 10 minutes for the indicated period.

### Fixation and immunostaining

Appropriately staged embryos were collected on a mesh, rinsed with Milli-Q water, and dechorionated using 70%(v/v) commercial bleach until most of the embryos had lost the chorion (confirmed visually). Embryos were fixed in a 1:1 mix of heptane and 4% paraformaldehyde in 1X phosphate buffered saline (1X PBS) at room temperature with nutation for 20 minutes. After removing the aqueous layer, an equal volume of 100% methanol was added, and embryos were devitellinized by vigorous manual shaking for 2–3 minutes. Fixed embryos were stored in 100% methanol at –20 °C.

For staining, embryos were rehydrated in 1X PBS, then washed, permeabilized and blocked twice for 15 minutes in PBTB (1X PBS, 0.2% Triton X-100, 1 mg/mL bovine serum albumin (Sigma-Aldrich, A9418). Embryos were then incubated with primary antibodies diluted in PBTB overnight at 4 °C. The next day, embryos were washed (4 × 15 min, PBTB) and incubated overnight at 4 °C with fluorophore-conjugated secondary antibodies and DAPI diluted in PBTB containing 4% normal goat serum (Jackson Immunoresearch, 005-000-121). On the third day, embryos were washed (4 × 15 min in 1X PBS) and mounted in Vectashield (Vector laboratories, H1000) for imaging. Primary antibodies used were rat anti-E-cadherin (1:20; DSHB, DCAD2), chicken anti-Vasa (1:800) (Repouliou et al., 2025) and rabbit anti-GFP conjugated to AlexaFluor 488 (1:1000; Invitrogen, A-21311). Secondary antibody used was goat anti-rat (1:200) conjugated to AlexaFluor 633 (Invitrogen, A-21094), goat anti-chicken (1:200) conjugated to AlexaFluor 568 (Invitrogen, A-11041). DNA was stained with DAPI (Sigma-Aldrich, D9542) at 1:2000 dilution of a 10 mg/mL stock.

### Image Analysis and Quantification

Live cell tracking was performed using the built-in Mastodon plugin in Fiji/ImageJ (71). Cell position relative to the gut lumen was determined manually for each time point. Fluorescence intensities were quantified by drawing manual regions of interest and measuring mean gray values using basic tools in Fiji/ImageJ (71). Germ cell positions, gut lumen boundary, and the center of the lumen were marked using the built-in Cell Counter function in Fiji/ImageJ (71). Germ cells were classified as “Out” if their distance from the lumen center exceeded that of the lumen boundary; otherwise, they were classified as “In”.

### In silico model of transepithelial migration

A cross-sectional slice of the epithelial midgut was modelled as a two-dimensional ring formed by 20 epithelial cells. The epithelial ring is embedded in a radial chemical field representing the spatial concentration of the chemoattractant. The magnitude of the chemical concentration increases linearly with the radius, and the minimum coincides with the center of the epithelial ring. At t=0, the germ cell begins slightly off-center, with its initial angle randomly chosen between 0 and 360 degrees, and then migrates outward toward regions of higher chemoattractant concentration.

Our two-dimensional model of the epithelial ring and the germ cells was formulated in the Cellular Potts Model (CPM) framework (38–40). The cells in a CPM are represented as a contiguous collection of pixels. Pairwise interactions between pixels at the cell boundaries have an associated energy cost, 𝐽. Given two pixels 𝑎 and 𝑏, we use the value of 𝐽_𝑎,𝑏_ to reflect the adhesion energy of the cells to which the two interacting boundary pixels belong. Our model assumes that adhesion energy increases with the number of adherens junctions formed, and that the number of adherens junctions is in turn a function of the E-cadherin concentration. The maximum possible number of junctions between two cells is thus determined by the surface E-cadherin concentration of the cell that has fewer such molecules. As a result of our assumptions, if σ(𝑎), σ(𝑏) are the indices of the cells to which pixels 𝑎 and 𝑏 belong, and 𝐶(σ(𝑎)) and 𝐶(σ(𝑏)) are the E-cadherin concentrations of the two cells, then the interaction energy term is 𝐽_𝑎,𝑏_ = *m*𝑖𝑛(𝐶(σ(𝑎)), 𝐶(σ(𝑏))). In our work, we have assumed that all epithelial cells have E-cadherin concentration equal to 𝐶_*E*_, and that all germ cells have E-cadherin concentration equal to 𝐶_*G*_. The energy function of the system of pixels is given by the following equation:

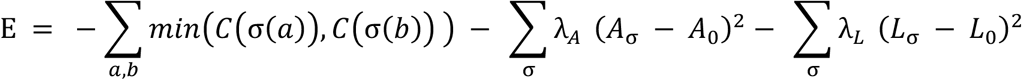

The λ_*A*_ and λ_𝐿_ terms determine the magnitude of the energy cost required for a cell area and perimeter to deviate from the steady state values *A*_0_ and 𝐿_0_ respectively. The system is simulated using the metropolis algorithm (45), where the probability of a pixel flip is related to the change in the magnitude of the energy of the system as a result of that specific pixel flip. Under appropriate energy conditions, pixels at the boundary may flip to the cell state of one of their neighbors that has a different cell state. At each iteration of the simulation algorithm, one such potential pixel flip is considered, and the pixel is flipped if the resulting system has lower energy than the current state. If the pixel flip would not reduce the energy of the system, the pixel flips with probability 𝑝_𝑓𝑙𝑖𝑝_ = 𝑒𝑥𝑝(−Δ*E*/𝑇), where Δ*E* is the change in energy due to the pixel flip and 𝑇 is the temperature. The temperature term 𝑇 represents the extent of stochasticity (noise) in the pixel dynamics, with higher 𝑇 increasing the probability of pixel flips that increase the energy of the system. Chemotaxis is implemented within the CPM framework by biasing pixel flips that aid cell movement in the direction of the gradient of chemoattractant. In the presence of the attractant whose value at any pixel in 2D space is 𝑀_𝑎,𝑏_, the change in energy due to a pixel flip, where a pixel at (𝑎, 𝑏) is copying the state of a pixel at (𝑎’, 𝑏’) becomes Δ*E*^6^ = Δ*E* − λ_𝐶_ (𝑀_𝑎’,𝑏’_ − 𝑀_𝑎,𝑏_) (40). This additional term favors the pixel flip when the chemical field value at (𝑎’, 𝑏’) is higher than (𝑎, 𝑏). To prevent the epithelial ring from collapsing and forming a cell aggregate without a lumen, we model the lumen as one large cell with a volume constraint. In addition, we mechanically couple the epithelial cells by connecting the centroids of the adjacent cells with springs using the FocalPointPlasticity plugin of CompuCell3D (72). The key parameters that we explored in our work are 𝐶_*G*_, 𝐶_*E*_ and λ_𝑀_. The CompuCell3D implementation of our model can be found at https://github.com/boyonpointe/GermCellTransepithelialMigration with commit ID 7ce2534.

## Acknowledgements

We thank members of the Extavour Lab for helpful discussions, in particular Dr. Emily Rivard and Dr. Petra Kovacikova for their comments on the manuscript. We thank Prof. James Glazier for helpful discussions regarding CompuCell3D, the Drosophila Genomics Research Center (NIH Grant 2P40OD010949) and Dr. Xiaona Tang for their help in making the E-cadherin-mScarlet CRISPR flies. We thank Ruth Lehmann for sharing fly lines.

## Data Availability

Scripts for Compucell3D simulations and image analysis are available at https://github.com/boyonpointe/GermCellTransepithelialMigration with commit ID 7ce2534.

## Author Contributions

CK conceived of the study, designed and performed all computational experiments, performed data analysis and interpretation, and wrote the first draft of the manuscript. SG conceived of the study, designed and performed wet lab experiments, and reviewed and edited the manuscript. CGE obtained funding for the study, supervised its execution, and reviewed and edited the manuscript.

## Funding

This study was supported by a postdoctoral fellowship to CK through the NSF-Simons Center for Mathematical and Statistical Analysis of Biology at Harvard (award number DMS-1764269), the Harvard Quantitative Biology Initiative, and by funds from Harvard University and the Howard Hughes Medical Institute (HHMI). CGE is an HHMI Investigator.

## Conflicts of Interest

The authors declare no conflicts of interest.

## Notes

### Competing Interest Statement

The authors have declared no competing interest.

### Summary of Updates

We have performed additional new computational and in vivo experiments, the results of which are included in the manuscript as additional supplemental figures and movies. The manuscript has been edited to describe the results of these new experiments, which strengthen and expand our original findings.

