## Supplemental Information for "Heterotypic intercellular adhesion tunes efficiency of cell-on-cell migration"

**This PDF file includes:**

Supporting Methods
Figures S1 to S7
Legends for Movies S1 to S7
SI References

### Supporting Methods

#### Generation of endogenous *shotgun–mScarlet* allele
We generated an endogenous C-terminal mScarlet fusion of *shotgun* (*shg*) in *Drosophila melanogaster* using scarless CRISPR/Cas9-mediated homology-directed repair ***(1)***.

#### Guide RNA plasmid
We designed the guide RNA (gRNA) sequence close to the end of the coding region of the shotgun gene. We used the sequence CGCAAATTGGCCGACATGTACGG, which maps to chr2R:21050864. The last three nucleotides (CGG) correspond to the PAM sequence. We altered CGG to CCG in the CRISPR donor plasmid using site-directed mutagenesis to prevent Cas9-mediated cleavage. To generate the guide RNA plasmid, we linearized the pU6-3-gRNA plasmid (DGRC #1362) using the primers CCGACATGTAgttttagagctagaaatagcaag and CCAATTTGCGcgaagttaaattgaaaatagg. The uppercase bases correspond to the primer overhangs matching the guide RNA sequence. Upon ligation of the linearized plasmid, we inserted the guide RNA sequence at the appropriate site in the template plasmid to generate the final guide RNA construct.

#### CRISPR donor plasmid
We constructed the donor plasmid using the pHD-ScarlessDsRed (Drosophila Genomics Resource Center Stock 1364; https://dgrc.bio.indiana.edu//stock/1364; RRID:DGRC_1364) template plasmid, which contains a 3xP3-DsRed selection cassette for screening HDR events. We designed the construct to insert the mScarlet coding sequence in-frame immediately downstream of the *shotgun* (*shg*) coding sequence. To minimize steric hindrance during folding of DE-cadherin and mScarlet, we added a 15 bp SG linker encoding GGSGS upstream of the mScarlet start codon.

We amplified the left (1003 bp) and right (1000 bp) homology arms from genomic DNA. To ensure seamless excision of the selection cassette, we inserted a 280 bp genomic sequence downstream of the shotgun stop codon and positioned it immediately upstream of the 3xP3-DsRed cassette. We selected this 280 bp fragment to include a TTAA site at its 3′ end so that PBac-mediated excision of the cassette would restore the endogenous genomic sequence without introducing an ectopic TTAA site.

We assembled the final donor plasmid in three steps. In step 1, we linearized the pHD-ScarlessDsRed template plasmid using round-the-horn PCR and inserted three fragments: (i) the left homology arm (1003 bp), (ii) the mScarlet coding sequence (693 bp), and (iii) the 280 bp sequence ending with TTAA. We amplified all three fragments from genomic DNA of a fly line carrying squash tagged with mScarlet (*w[*] TI{TI}sqh[mScarlet-I]; MKRS/TM6B, Tb[1]*, Bloomington Drosophila Stock Center (BDSC) #94929). We designed primers to introduce ~25 bp overlaps between adjacent fragments and with the linearized backbone and assembled the fragments using Takara In-Fusion Snap Assembly. In step 2, we linearized the partially assembled plasmid using round-the-horn PCR and inserted the right homology arm, which we amplified from genomic DNA of BDSC #94929 with ~25 bp overlaps to the backbone. We assembled the insert and backbone using Takara In-Fusion Snap Assembly. In step 3, we mutated the PAM site in the left homology arm using the NEB Q5 Site-Directed Mutagenesis kit to prevent Cas9-mediated cleavage and generate the final donor plasmid.

#### Embryo injection and screening
We injected the sgRNA and donor plasmids into embryos of genotype *yw;;nos-Cas9(III-attP2*, which express Cas9 in the germ line. BestGene (Chino Hills, CA) performed the injections. We crossed G0 adults to a white-eyed second chromosome balancer line (*yw; If/Cyo; +/+*). We screened progeny for DsRed fluorescence in the eyes to identify candidate HDR events. We confirmed the insertion by PCR using a forward primer binding upstream of the left homology arm and a reverse primer binding within the mScarlet sequence.

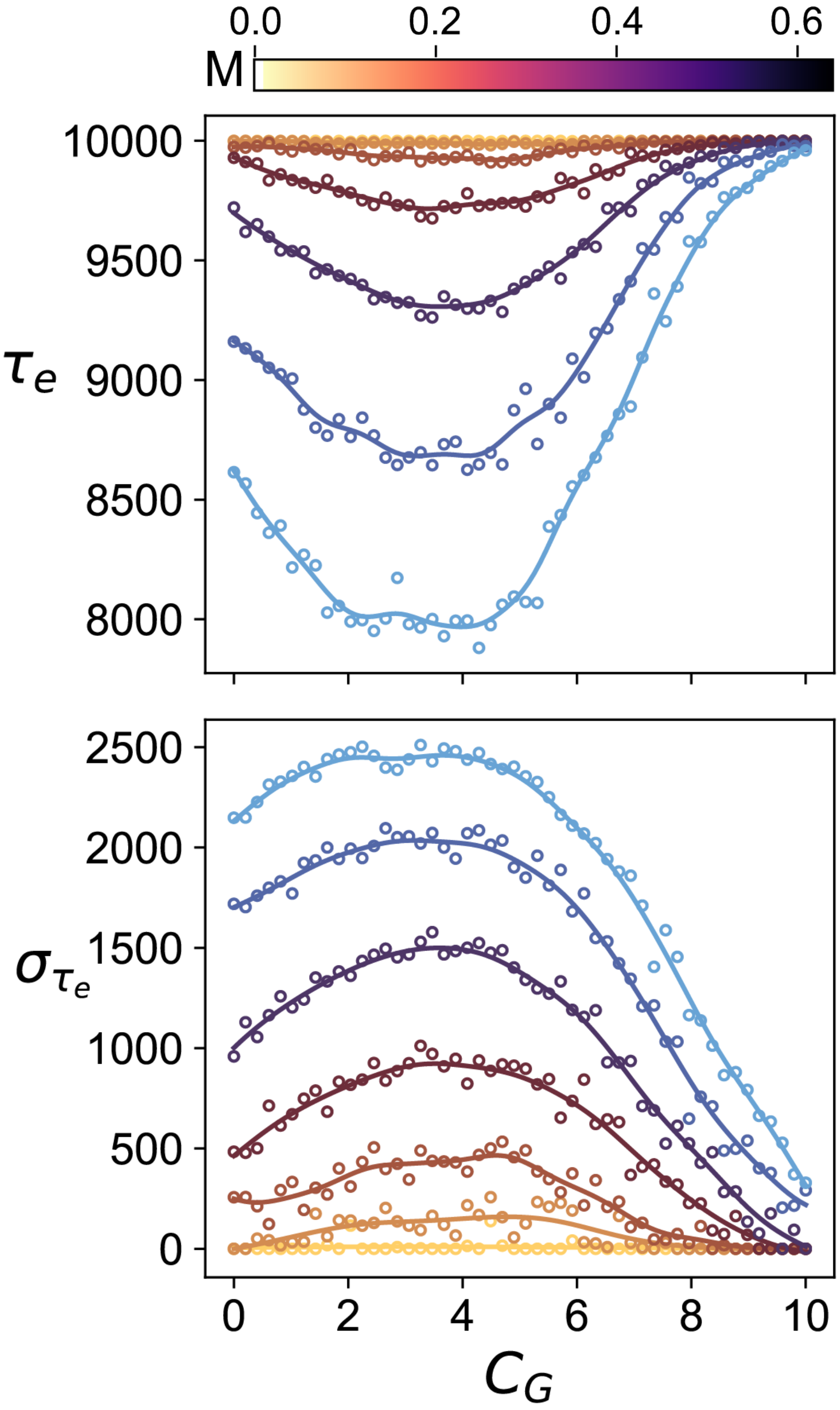


**Fig. S1. The mean and the variance of $\tau_e$ exhibit converse trends.** The figure shows the mean $\boldsymbol{\tau_e}$ (top) and standard deviation $\boldsymbol{\mu_{\tau_e}}$ (bottom) as a function of germ cell E-cadherin concentration $\boldsymbol{C_G}$ for the case when $\boldsymbol{C_E}$ = 10. The peak in the standard deviation as a function of $\boldsymbol{C_G}$ coincides with the minima in the case of time to exit. The differently colored curves correspond to different strengths of the chemoattractant pull $\boldsymbol{\lambda_M}$.

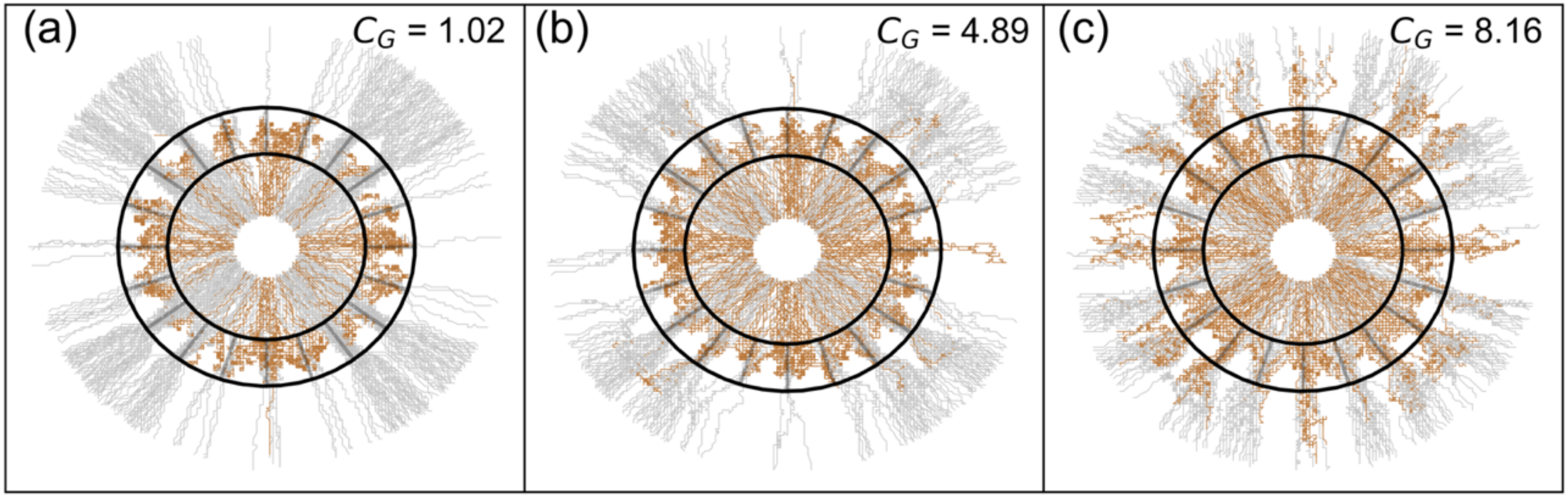


**Fig. S2. Insufficient and excessive heterotypic adhesion represent two distinct failure modes in transepithelial migration.** Figure showing trajectories at one value lower ( $C_G$ = 1.02) and one value higher ($C_G$ = 8.16) than the condition shown in Figure 3c ($C_G$ = 4.89). As before, gray trajectories correspond to cells that successfully exited, while brown trajectories correspond to cells that failed to exit. At lower heterotypic adhesion ($C_G$ = 1.02), it is evident that most failed cells have trajectories ending more apically, indicating they could not breach the epithelial barrier. Conversely, at very strong heterotypic adhesion ($C_G$ = 8.16), failed cells stall more basally, with much of the cell body remaining outside the epithelial barrier, as reflected by the numerous brown trajectories ending outside the ring. Note that since the trajectories track the position of the center of mass, a center of mass outside the ring does not necessarily indicate that the entire cell body has exited.

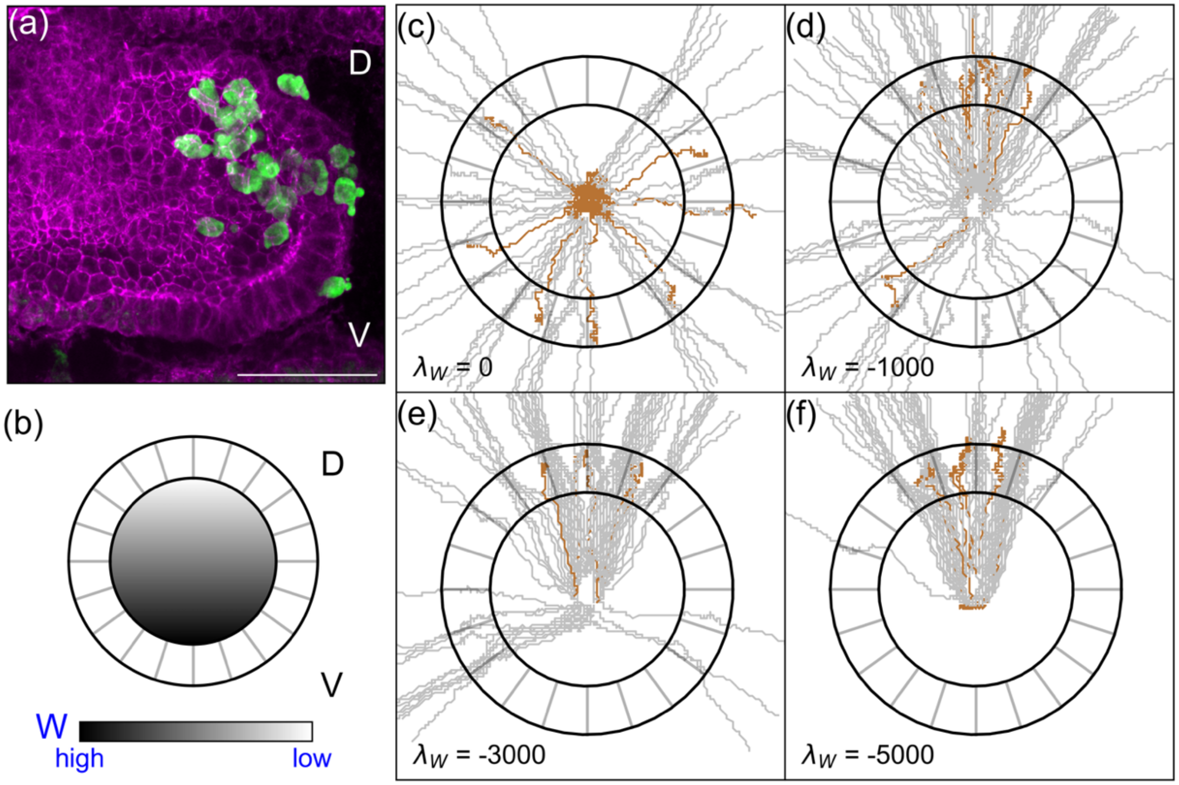


**Fig. S3. Wun/Wun2 repulsion biases germ cell dispersal toward the dorsal side during transepithelial migration.** (a) Fluorescence micrograph showing germ cells (green) exiting the midgut preferentially toward the dorsal (D) side. E-cadherin is shown in magenta. Scale bar, 50 µm. (b) Schematic of the simulated Wun/Wun2 (W) concentration field used in the model: W is highest near the ventral (V) pole and decreases linearly toward the dorsal pole, consistent with the known expression pattern of these lipid phosphate phosphatases along the dorsoventral (DV) axis **(2, 3)**. In the model, germ cells experience both a radially outward attractive cue driving exit from the midgut and a repulsive cue proportional to local W concentration. (c–f) Simulated germ cell trajectories during transepithelial migration for increasing strengths of Wun/Wun2-mediated repulsion ($\boldsymbol{\lambda_W}$ = 0, −1000, −3000, −5000). Individual cell tracks are shown in gray; the mean trajectory is highlighted in orange. Concentric circles indicate radial distance from the midgut center. With no repulsion ($\boldsymbol{\lambda_W}$ = 0), trajectories are distributed isotropically, whereas increasing $\boldsymbol{\lambda_W}$ progressively confines exit to the dorsal hemisphere, reproducing the dorsal bias observed experimentally.

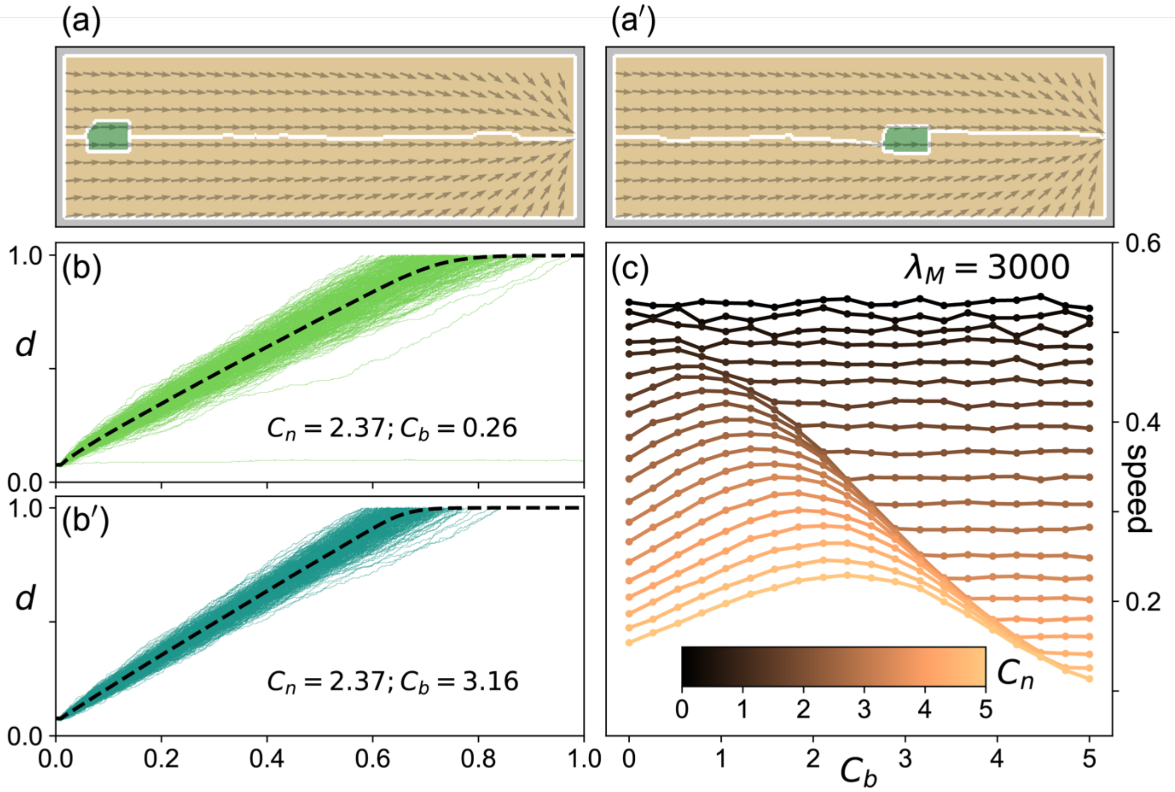


**Fig. S4. Optimal cell-substrate adhesion maximizes germ cell migration speed.** (a-a') Representative snapshots from CompuCell3D simulations of a single migratory cell (green) migrating through a channel formed by two substrate cells (tan) at two different time points during the simulation. Arrows indicate the local chemotactic gradient field and white outlines mark cell boundaries. (b-b') $\boldsymbol{C_n}$ and $\boldsymbol{C_b}$ are the E-cadherin concentration in the substrate cells and the migratory cell respectively. (b),(b') show the normalized cell trajectories lower , $\boldsymbol{C_b}$= 0.26 and higher value, $\boldsymbol{C_b}$ = 3.16 of migratory cell adhesion for the same value of substrate cell adhesion, $\boldsymbol{C_n}$ = 2.37. Individual simulation trajectories are shown as colored lines and the mean trajectory as a dashed black line. The normalized displacement *d* (0.0-1.0) is plotted against normalized time (0.0–1.0). (c) Mean migration speed as a function of substrate cadherin concentration $\boldsymbol{C_n}$ and migratory cell cadherin concentration $\boldsymbol{C_b}$ at fixed strength of chemoattractant $\boldsymbol{\lambda_M}$= 3000. For low $\boldsymbol{C_n}$ (dark curves) where there is no adhesion between the substrate cells speed is largely insensitive to $\boldsymbol{C_b}$. For intermediate-to-high $\boldsymbol{C_n}$ (lighter curves), speed exhibits a clear peak at intermediate $\boldsymbol{C_b}$, demonstrating that an optimal substrate–cell adhesion strength maximizes migration speed. Excessive adhesion stalls the cell, while insufficient adhesion fails to provide the traction needed for efficient migration.

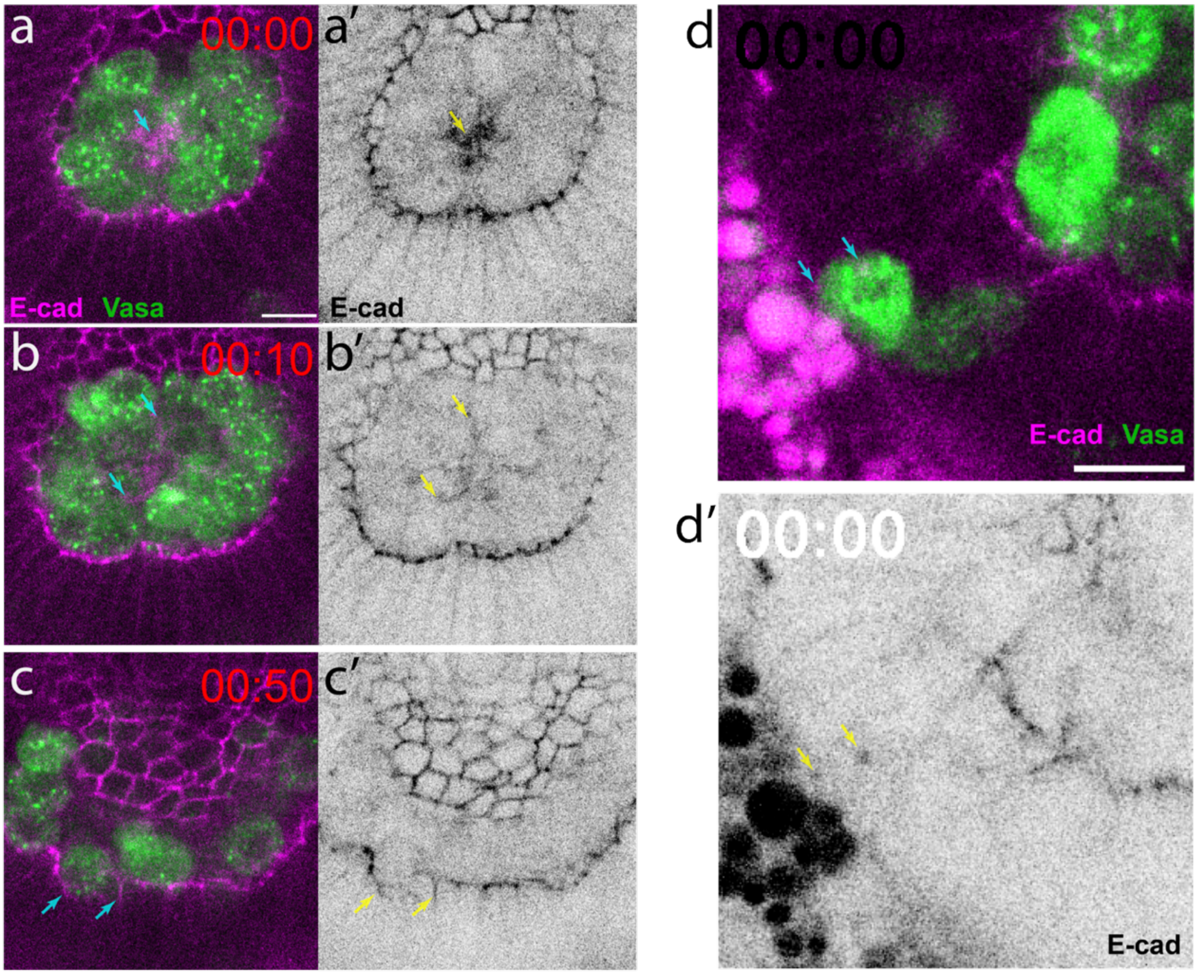


**Fig. S5. Germ cell transepithelial migration and exit from the gut.** Selected frames from (a-c') SI Video 6, and (d,d') SI Video 7. Panels a,b,c and d show localization (gray arrows) of E-cadherin-mScarlet with respect to germ cells expressing Vasa-GFP in an embryo 5.5 hours after egg-layingat different timepoints of the time lapse video. For clarity, panels a',b'c', and d' show corresponding localization (yellow arrows) of only E-cadherin-mScarlet. Scale bars are 10μm.

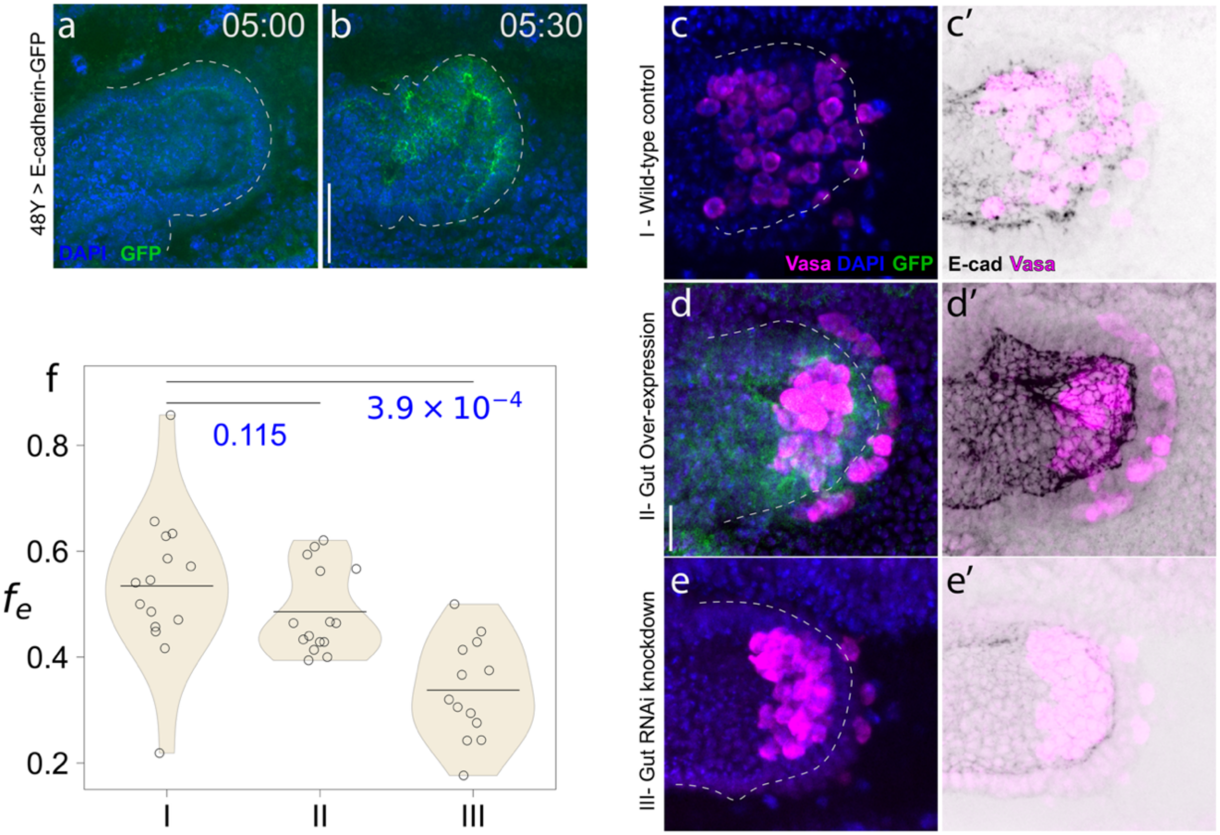


**Fig. S6. Epithelial perturbation of E-cadherin during transepithelial migration.** Zygotic expression of E-cadherin-GFP using the endodermal driver 48Y-Gal4 at (a) 5.0 hours and (b) 5.5 hours after egg laying (AEL). Germ cell positions and E-cadherin levels at 5.5 hours AEL in (c,c') I: Wild-type control (48Y-Gal4/Oregon R , n = 15) (d,d') **II**: Gut E-cadherin over-expression (48Y-Gal4>E-cadherin=GFP, n=15), and (e,e') **III**: Gut E-cadherin knockdown (48Y-Gal4>E-cadherin-RNAi, n=13).(f) The fractions of germ cells that have successfully exited the gut ($f_e$) is shown for the three different cases. There is no statistically significant difference observed between the wild type (I) and the embryos with gut specific overexpression of E-cadherin (II). In the case of gut specific E-cadherin knockdown (III) we see statistically significant delay compared to the wild type (I). Two sample Mann-Whitney U test was used to compute the p values that are shown in panel (f) for pairwise comparisons of distributions. Dashed line indicates the basal surface of the gut epithelium. Scale bar represents (a,b) 50μm and (c-e') 20μm. Images are maximum intensity projections of (a,b) 10μm and (c-e') 20μm thickness.

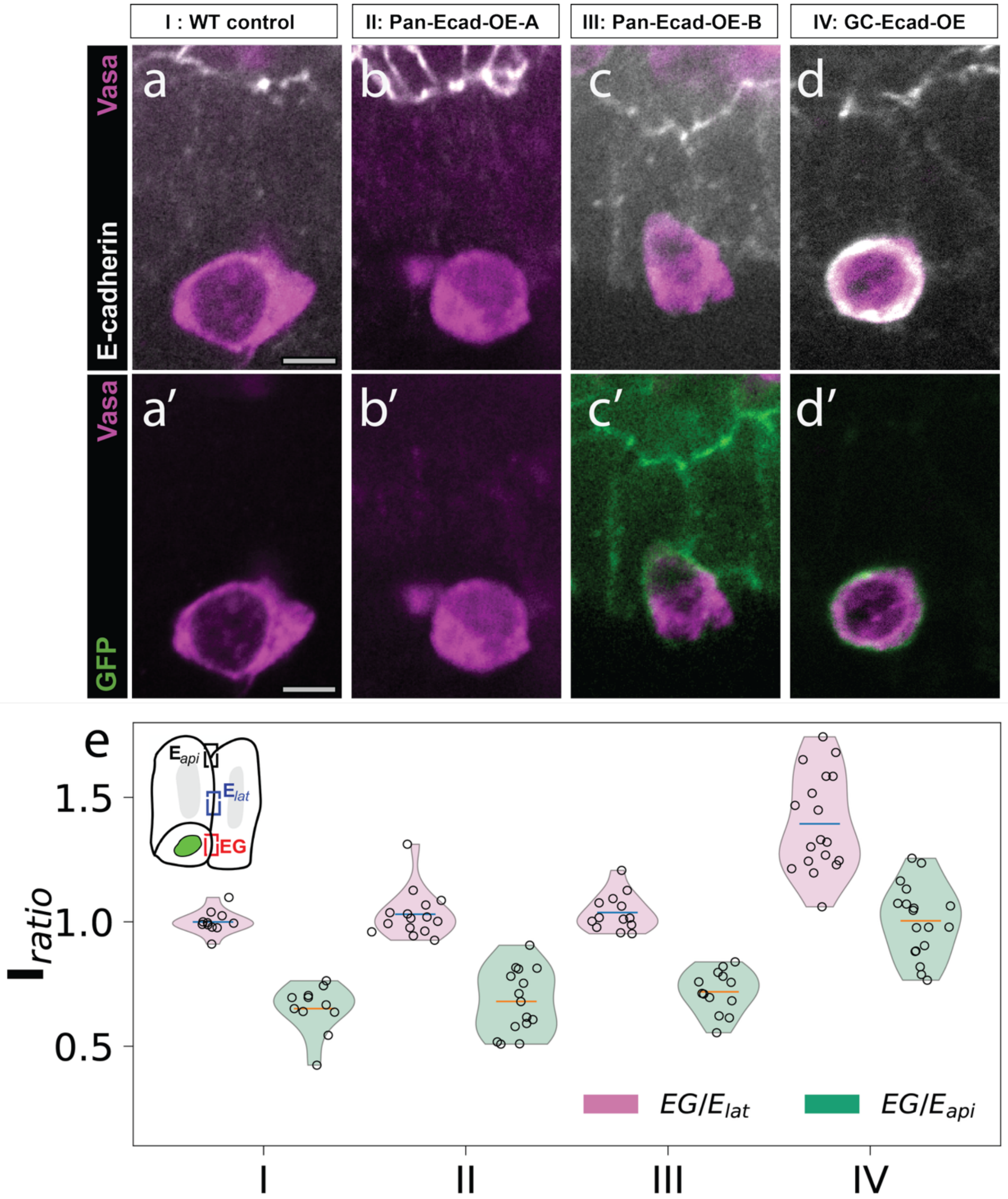


**Fig. S7. E-cadherin levels during heterotypic adhesion.** Cross-sectional view of a single germ cell in the gut epithelia during TEM in similar genotypes as Figure 4a, (a-a') I: Wild-type control (n = 11), (b-b') II: Pan-embryonic overexpression of E-cadherin (n=15), (c-c') III: Pan-embryonic over-expression of E-cadherin-GFP (n=12) and (d-d') IV: Germ cell-specific over-expression of E-cadherin-mClover2 (n=18). Scale bar represents 5µm. (e) The E-cadherin amount at the surface of the germ cell (EG) relative to E-cadherin in the lateral junction (pink) and apical junction (green) of the epithelial cells are shown for several junctions spanning multiple embryo samples for the four different experimental groups. The germ cell-specific over expression condition (IV) shows significantly higher relative germ cell E-cadherin than the wild type (I). In contrast, the two different pan-embryonic overexpression conditions only show a slight increase in germ cell E-cadherin expression relative to the epithelial E-cadherin expression.

### SI Movie Legends

**Movie S1 (separate file). Transepithelial migration of germ cells.** Dorsal cross-sectional view of the gut during TEM in a *nos-LifeAct-tdTomato* embryo. Maximum intensity projection of 15 planes spaced at 2 µm. Scale bar is 20 µm.

**Movie S2 (separate file). Transepithelial migration of germ cells: higher magnification 1**. Dorsal cross-sectional view of the gut during TEM in a *nos-LifeAct-tdTomato* embryo. Maximum intensity projection of 20 planes spaced at 2 µm. Scale bar is 20 µm.

**Movie S3 (separate file). Transepithelial migration of germ cells: higher magnification 2.** Lateral cross-sectional view of the gut during TEM in a *nos-LifeAct-tdTomato* embryo. Maximum intensity projection of 20 planes spaced at 2 µm. Scale bar is 20 µm.

**Movie S4 (separate file). CompuCell3D simulation of a germ cell successfully exiting the midgut.** The video shows an instance of a germ cell successfully exiting the epithelial ring in our *in silico* model simulated in CompuCell3D. The parameter values used here are $\boldsymbol{\lambda_M = 4000, C_G = 5.0, C_E = 10.0}$.

**Movie S5 (separate file). CompuCell3D simulation of a germ cell failing to exit the midgut**. The video shows an instance of a germ cell failing to exit the epithelial ring in our *in silico* model simulated in CompuCell3D. The parameter values used here are $\boldsymbol{\lambda_M = 4000, C_G = 1.0, C_E = 10.0}$.

**Movie S6 (separate file). Transepithelial migration of germ cells.** Dorsal cross-sectional of view of the gut during TEM in live embryos expressing Vasa-GFP (green) and E-cadherin-mScarlet (magenta). Single plane imaged at an interval of 5 minutes.

**Movie S7 (separate file). Germ Cell exit from the gut.** Dorsal cross-sectional view of the basal surface of the gut during germ cells exit in embryo expressing Vasa-GFP (green) and E-cadherin-mScarlet (magenta). Single plane imaged at an interval of 1 minute.
